# Dual Pathways of Extracellular ATP Action in Cancer Cells: Purinergic Signaling–Driven Senescence and Macropinocytic ATP Internalization

**DOI:** 10.64898/2026.04.23.720363

**Authors:** Nicole Stone, Ryan Ward, Lindsey Bachmann, Subhodip Adhicary, Corinne M. Nielsen, Nikunj Mehta, Yunsheng Li, Haiyun Zhang, Jingwen Song, Sam Prinz, Stephen Chang, Dennis Roberts, Stephen Bergmeier, Pratik Shriwas, Xiaozhuo Chen

**Author notes:** These authors contributed equally (Nicole Stone, Ryan Ward). Correspondence authors: Correspondence to Xiaozhuo Chen, 109 Edison Biotechnology Institute Ohio University, 172 Water Tower Drive, the Ridges Athens, Ohio 45701, USA, Pratik Shriwas, Room 305, JNC, School of Chemical and Biotechnology, Sastra University, Thanjavur, Tamil Nadu 613401.

## Abstract

**Background:** Opportunistic nutrient uptake is a hallmark of cancer metabolism. Cancer cells upregulate macropinocytosis to acquire extracellular nutrients to support growth and stress adaptation. We previously showed that extracellular ATP (eATP) is internalized by macropinocytosis and promotes multiple cancer phenotypes. Here, we tested whether eATP uptake is prevalent across cancers and whether eATP also induces senescence through purinergic receptor (PR) signaling.

**Methods:** Intracellular ATP (iATP) levels were measured following eATP exposure across multiple cancer cell lines. eATP internalization was visualized in vitro and in vivo using a non-hydrolyzable fluorescent ATP analog together with high-molecular-weight dextran as a macropinocytosis marker. Senescence was quantified using three SA-β-galactosidase assays and flow cytometry. Pharmacologic inhibitors of macropinocytosis and purinergic receptors were used to define pathway dependence. Combination treatments with the glucose transporter inhibitor DRB18 and the senolytic navitoclax were evaluated for antiproliferative effects.

**Results:** eATP produced dose- and time-dependent increases in iATP across diverse cancer cell types. Imaging demonstrated widespread macropinocytic internalization of ATP in vitro and in tumor xenografts. eATP induced senescence in NSCLC cells, confirmed by multiple β-gal assays and flow cytometry. PR inhibition significantly reduced senescence, whereas macropinocytosis inhibition had minimal effect on senescence induction.

**Conclusions:** eATP acts through dual pathways in cancer cells: macropinocytic internalization that elevates iATP and PR signaling that drives senescence. Targeting metabolic uptake together with senolytic therapy may offer a novel anticancer strategy.

## Introduction

Extracellular ATP (eATP) is an important signaling molecule released from cancer cells and reaching concentration between 0.1–1 mM in the tumor microenvironment thereby infiltrating immune cells, and hypoxia-associated tumor cell lysis ^1,2^. At these elevated levels, eATP functions as a stress and danger signal and activates purinergic receptors (PR), particularly P2X7, to regulate multiple cancer phenotypes ^1–4^. Previous studies, including our own, have shown that tumor-level eATP induces epithelial–mesenchymal transition (EMT), cancer stem cell (CSC) formation, and drug resistance in non-small cell lung cancer (NSCLC) ^5–8^. These effects resemble those induced by TGF-β signaling, and PR activation has been linked to downstream pathways that promote EMT and CSC programs ^4,6^. Together, these findings suggest that eATP is a multifunctional regulator of tumor cell behavior and stress responses.

In addition to receptor-mediated signaling, cancer cells actively scavenge extracellular nutrients through macropinocytosis, a high-capacity endocytic process that engulfs extracellular fluid and macromolecules into large vesicles called macropinosomes ^9,10^. Macropinocytosis is enhanced in several oncogenic settings, including RAS-driven tumors, and contributes substantially to metabolic adaptation ^11,12^. We previously demonstrated that eATP is internalized via macropinocytosis in lung and breast cancer cells both in vitro and in vivo ^13–15^. This uptake increases intracellular ATP (iATP) levels and activates metabolic and signaling pathways that promote EMT, stemness, and drug resistance. Pharmacologic inhibition of macropinocytosis or genetic modulation of regulators such as STC1 reduces ATP internalization and reverses several of these cancer-promoting phenotypes ^7^. These findings suggest a major role for macropinocytic eATP uptake in tumor metabolism and therapy response.

Cellular senescence is a stress-responsive state characterized by enlarged morphology, stable cell-cycle arrest, and a senescence-associated secretory phenotype (SASP) ^5,16–18^. Although historically considered an anticancer mechanism was targeted by our glucose transporter inhibitor WZB117 ^19–21^, senescence is now recognized as context-dependent and often cancer-promoting ^10,24–25^. Senescent tumor cells can remodel the microenvironment through SASP, enhance malignant traits in neighboring cells, contribute to therapy resistance, and in some cancers reenter the cell cycle with increased stem-like properties ^5,26–27^. Senescence is commonly detected by senescence-associated β-galactosidase (SA-β-Gal) activity driven by increased lysosomal biogenesis and galactosidase β1 (GLB1) expression ^28–29^, which correlates with poor patient survival. Senescent cells are therefore increasingly viewed as therapeutic targets, and senolytic agents, a class of drugs designed or identified to selectively eliminate senescent cells have emerged as a new anticancer strategy ^30–31^.

Based on the established roles of eATP in PR signaling and macropinocytic uptake, we hypothesized that eATP exerts dual but distinct effects in cancer cells: macropinocytosis-mediated ATP internalization and PR-mediated senescence induction. In this study, we tested the prevalence of eATP macropinocytosis across multiple cancer types and determined the signaling mechanism of eATP-induced senescence. Using quantitative ATP measurements, fluorescence imaging *in vitro* and *in vivo*, multiple SA-β-Gal assays, specific pathway inhibition for senescence and micropinocytosis, we concluded that eATP can induce dual and distinct mechanism in cancer cells.

## Materials and Methods

### Cell lines and cell culture reagents

All cell lines were purchased from American type tissue collection (ATCC) and were maintained in specific growth media supplemented with 10% FBS and 1% penicillin/streptomycin (Pen/strep). Cells were cultured in cell culture incubator at 37°C and 5% CO_2_. Cells were split using 0.25% trypsin-EDTA solution and washed with Phosphate buffered Saline (PBS) as and when required. NL-20 was a non-tumorigenic lung cell line used as a negative control for macropinocytosis. Table 1 summarizes the cell lines with growth medium. A549 STC1KO cells were generated as previously described ^7^. IPA3 was dissolved in DMSO and control condition 0.1% DMSO was used.

**Table 1.**
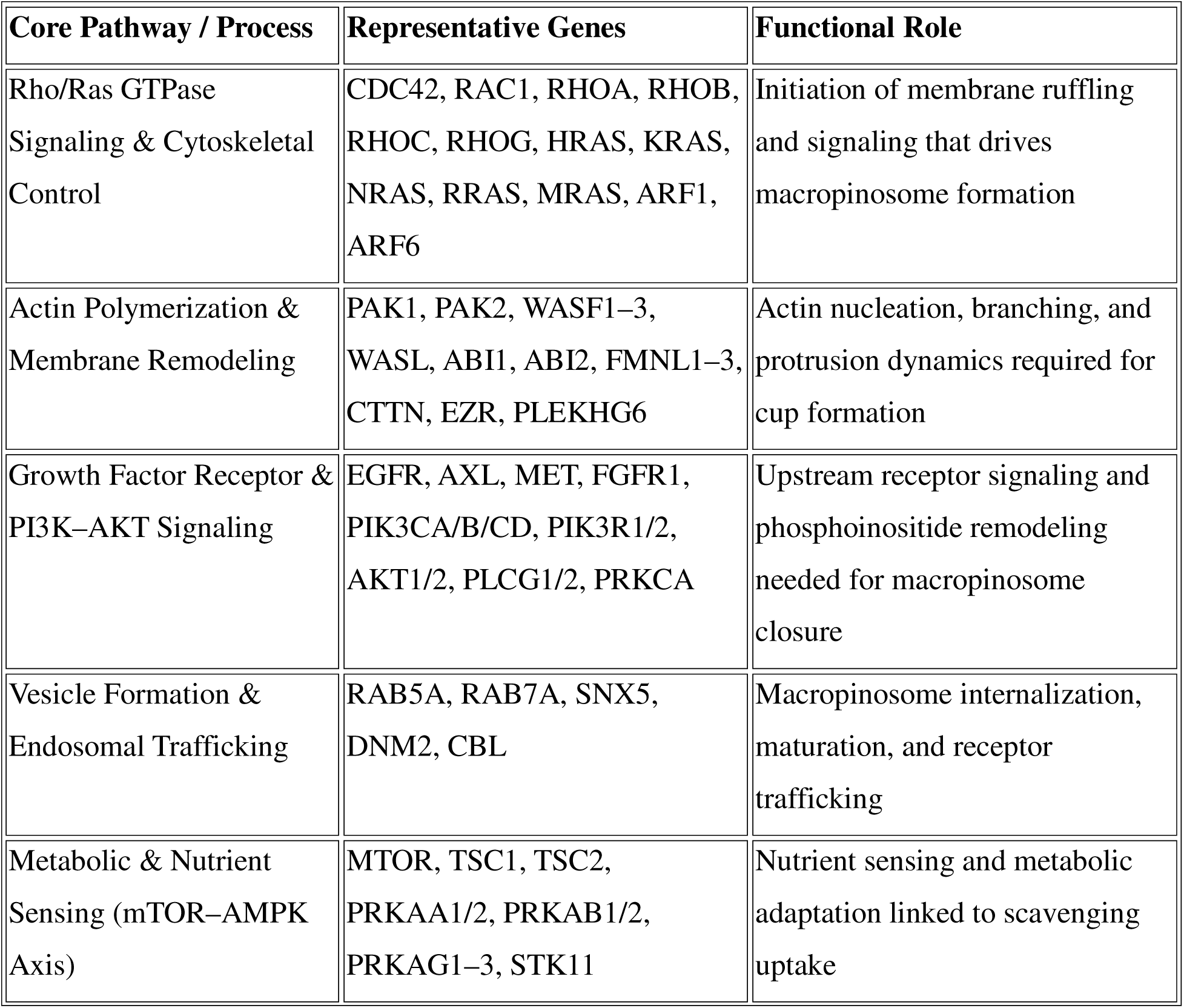
Cell lines and cell culture media.

**Table 1.**
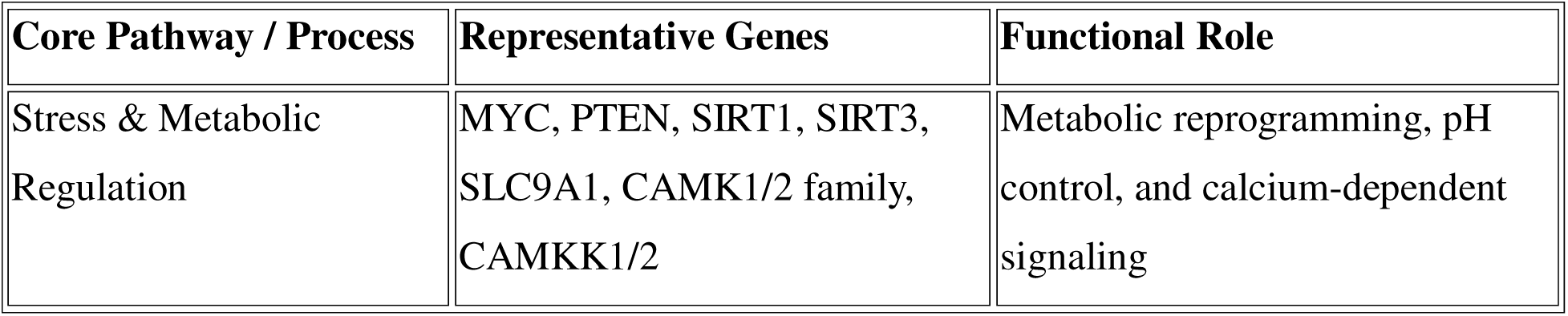
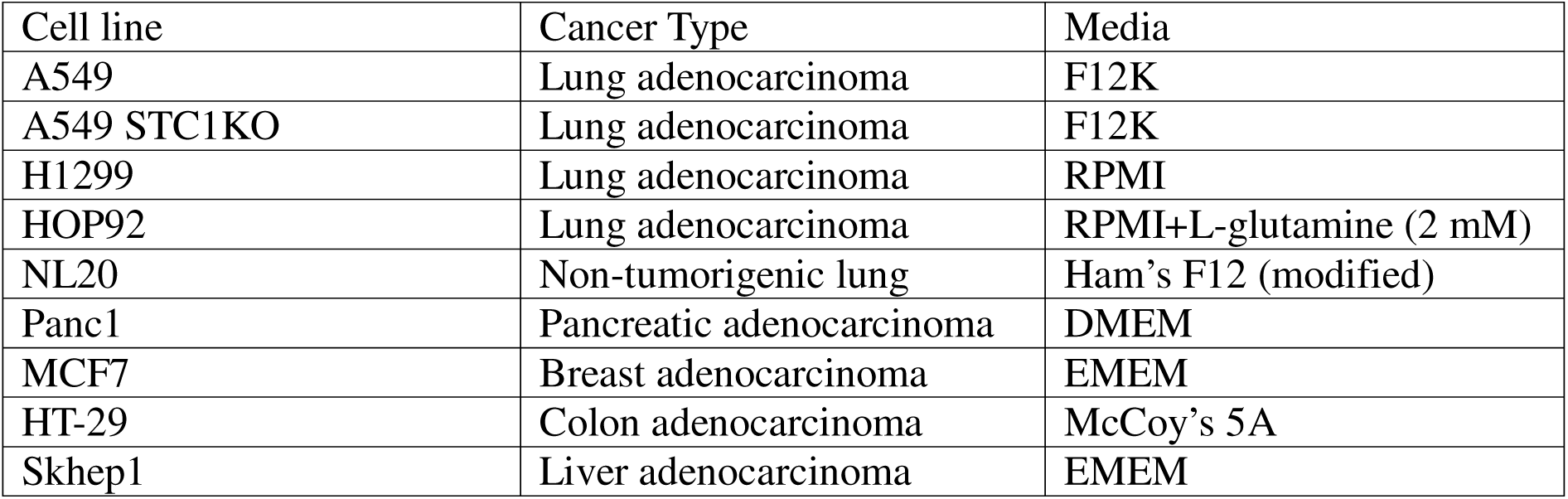
Macropinocytosis Signature Genes Grouped by Core Functional Pathways.

### Compounds inhibitors and other chemicals

Adenosine 5-triphosphate disodium salt hydrate (A2383), Resazurin sodium salt (R7017), 5-(N-Ethyl-N-isopropyl)amiloride (EIPA) (A3085), IPA3 (I2285), Navitoclax (AMBH2D6F59B5) and 200 mM glutamine (59202C) were purchased from Sigma Aldrich. DRB18 was generously supplied by department chemistry, Ohio University. Dulbecco’s Modified eagle Medium (DMEM) high glucose (ATCC-30-2002), Eagle’s Minimum Essential Medium (EMEM-ATCC 30-2003), RPMI (ATCC-30-2001), F12 (ATCC-30-2007), Ham’s F12 modified (ATCC-30-2004), McCoy’s 5A medium (ATCC-30-2007), 0.25% Trypsin-EDTA Solution, 1X (ATCC-30-2101), Fetal bovine serum (ATCC-30-2020), and Penicillin-Streptomycin Solution (ATCC-30-2300) were purchased from ATCC. For fluorescence microscopy, NHF-ATP was obtained from Jena Bioscience (NU-810-488), ProLong^TM^ Gold Antifade Mountant with DAPI from Thermo Fisher and high molecular weight fluorescent TMR-dextran (HMWFD) 70,000 Daltons MW, Neutral (no electric charge) from Invitrogen (D818).

### Intracellular ATP assay (short and long duration)

Intracellular ATP assay was performed as previously described ^19,30^. Briefly, 20000-50000 cells were seeded in black 96 well plate. The cells were allowed to adhere and grow for 24 hours. The cells were then washed with PBS twice and then treated with fresh media containing appropriate amount of ATP (0, 0.25, 0.5, 0.75 mM); for short assay 0, 30, 60, 90 and 120 minutes and longer assay 2,3 and 4 hours. All the treatments were timed such that all wells could be read at the same time. Finally, after desired incubation cells were washed with PBS twice and then 30 µl of mammalian cell lysis solution was added to each well. The plates were shaken at ∼700 rpm for 5 minutes. Next, 30 µl of substrate solution was added and plates were shaken in dark for 5 minutes at ∼700 rpm and then incubated in dark for another 10 minutes without shaking.

Luminescence reading was taken using biotek plate reader with gain setting of 150. To determine the effect of macropinocytosis inhibition, IPA3 (0, 10, 30, 50, and 100mM) and 0.5mM ATP were added to the wells. After 2 hours of treatment, the cell media was removed, and cells were washed with PBS twice. The remainder protocol was similar to as described above for eATP dosage study. The experiments were performed with 4 to 6 technical replicates for each condition and the experiment was repeated three times for biological replicates.

### Fluorescence microscopy *in vitro*

Fluorescence microscopy was performed as previously described ^31^. Non-hydrolysable fluorescent ATP (2’/3’-O-(2-Aminoethyl-carbamoyl)-Adenosine-5’-[(β,γ)-imido] triphosphate) exhibiting positivity green supplemented with 10mM regular ATP was used as ATP treatment. Cells (60,000-100,000) were seeded on coverslips in 24 well plates and incubated with serum containing media overnight. Next day, cells were washed twice with PBS and then incubated for 40 minutes with serum free media containing 0.75 mg/ml HMWFD along with 10 µM NHF-ATP and 10 mM regular ATP was added. After incubation, cells were washed three times with PBS and fixed with 4% paraformaldehyde for 15 minutes. Cells were then washed, and coverslips were removed from 24 well plates and mounted on clean microscope slides containing 10 µl DAPI. Negative control (DAPI only), NHF-ATP with DAPI, and HMWFD with DAPI. NHF-ATP with DAPI and HMWF-Dextran with DAPI images were combined using a Carl Zeiss Microscopy Axio Observer 7 Microscope. For A549 vs A549 STC1KO vs NL20 cells comparison, images were quantified using ImageJ software.

### Animal study

All animal studies protocol complied with with US regulations for animal usage in laboratory settings as well as IACUC policy of Ohio University, USA.

*In vivo* ATP internalization was performed as previously described ^14,31^. 3 to 4 weeks old male *NU/J* nude mice were purchased from The Jackson Laboratory and were fed with the Irradiated Teklad Global 19% protein rodent diet (Harlan Laboratories). Mice were allowed for one week of acclimatization and then 1×10^6^ – 5×10^6^ cells were injected subcutaneously into the right flank. Tumors were allowed reach palpable size of 200 – 500 mm^3^ and then injected with three conditions for each cell line: control (DMEM only), 8 mg/mL HMWFD along with or without (100 μmol/L) NHF-ATP in DMEM. The total injection volume was 50 μl using 1 CC syringes with 27G needles. Three tumors were injected for each condition. After injection, the mice were euthanized, and tumors were removed and then frozen on OCT with the entire process from injection to OCT freezing lasting about 7-8 minutes. Serial sections from tumor tissues of 10 µm thickness were collected throughout the tumor volume to account for intratumoral spatial heterogeneity. Collected sections were fixed in 95% ethanol at −20 °C for 5 minutes, washed in PBS and mounted using antifade medium containing DAPI. Images were acquired using a Nikon NiU epifluorescence microscope; and acquisition parameters were held constant across experimental groups. Image processing and analysis were performed using Nikon NIS-Elements software.

### Senescence study

Senescence studies were performed as previously described ^6^. Three senescence Associated β-Galactosidase Chromogenic Assays were performed to quantitively validate the effect of eATP in senescence induction ^32^.

#### A. Manual counting method

To selectively stain for senescent cells, a chromogenic assay using X-gal was utilized to detect and measure GLB1 activity. X-gal, a lactose analogue, is the most commonly used SA β-galactosidase substrate. By the action of GLB1, X-gal is cleaved into galactose and 5-bromo-4-chloro-3-hydroxyindole and then dimerizes to form a blue-colored compound.

After ATP and other treatments, A549 cells in 96-well or 48-well plates cells, depending upon assay types, were washed with PBS following by fixation with 2% formaldehyde and 0.2% glutaraldehyde in PBS for five minutes at room temperature. After fixation, cells are washed twice with PBS and then treated with a staining solution containing 1 mg/mL of X-gal and buffered to a pH of 6.0 with a citrate-phosphate buffer overnight for 12 to 16 hours. Cells were then washed twice with PBS, once with methanol, allowed to dry, and observed under a ZEISS Axio Observer inverted platform microscope on a bright field setting. For quantification of senescent cells, standard representative samples of each treatment well are observed and manually counted under a microscope.

### Additional methods

To validate the manual counting results and increase the reliability of SA β-Gal detection, GLB1 activity in A549 and H1299 cells was also measured by two additional methods:

The cell culture and cell treatment parts of these two assays were the same as the manual counting method described above, except that, for H1299 cells, 48-well plates were used for better monolayer formation. A 48 well plate was seeded at 25% confluency 24 hours before assay began for acclimatization. The media was removed and washed with PBS. Next treatment media were added (0.5mM ATP with DMSO at 5% by volume, paclitaxel in RMPI at 101.4 nM (the IC_50_ of paclitaxel) and DMSO at 5% by volume was used as positive control, and RMPI with 5% DMSO by volume as negative control).

#### B. X-gal fluorescence method

X-gal measured by a plate reader (BioTek Cytation 3) at 615nm

#### C. ONPG assay method

Cells were incubated for the desired time and then treatment media were removed and cells washed with PBS. The cells were then trypsinized and placed into 1.5 mL tubes and centrifuged at 12,000 g for 7 minutes. The supernatant was removed and then the pellet was resuspended in a 0.1 M phosphate buffer at a pH 6. The cells were then lysed via sonication, and the tubes were centrifuged again. The supernatant was retained and replated into a 48 well plate. 51 microliters of 2.2 μg/μL cell lysate in pH 6 phosphate buffer was added. Cells were then incubated at 37°C for ∼12 hours or longer. At the end of incubation, 87 μL of 1 M sodium carbonate was added, and the plate was read for absorbance at 420 nm with the plate reader.

For all three tests, each assay was repeated at least 3 times and samples of the same condition were 3-6 in each repeat (N=3-6).

### Flow Cytometry analysis

For a more quantitative measure of senescent cells, the CellEvent™ Senescence Green Flow Cytometry Assay Kit (ThermoFisher) were used according to manufacturer’s instructions. Briefly, following treatment, cells were washed with PBS and then trypsinized. Cells were then centrifuged at 1100 rpm at room temperature, the supernatant removed, and the cell pellets resuspended in PBS, then centrifuged and resuspended again in fixation solution containing 2% formaldehyde in PBS for ten minutes at room temperature. Following fixation, cells are centrifuged, supernatant removed and resuspended in 1% bovine serum albumin (BSA) solution in PBS and then centrifuged and resuspended in working solution supplied by the kit at 1000x dilution for 1-2 hours at 37°C in a non-CO_2_ incubator. After staining, cells are centrifuged and resuspended in 1% BSA twice and then analyzed using a BD FACSAria Cell Sorter Flow Cytometer with a laser at 488nm. This method led to revealing SA-β-Gal expressing population and eATP’s induction mechanisms.

### Bioinformatics analysis; Preparing for macropinocytosis score

The methods written below are for LUAD as an example. For other types of cancer, the names could be replaced with PAAD, LIHC, BRCA, PRAD, etc.

### Data Acquisition

RNA-seq gene expression (TPM) and clinical metadata for **TCGA-LUAD** (lung adenocarcinoma) and TCGA-PAAD (pancreatic adenocarcinoma) were downloaded from cBioPortal (TCGA Pan-Cancer Atlas, 2018).

### Macropinocytosis Gene Set

A curated list of **75 genes** involved in macropinocytosis was compiled based on literature, including genes regulating actin remodeling, small GTPases (e.g., RHO, RAC, CDC42 families), PI3K/AKT signaling, membrane trafficking (RABs, SNXs), and nutrient sensing (e.g., mTOR, AMPK).

### *In Silico* and Correlation Analyses

Transcriptomic and clinical survival data for TCGA-LUAD and TCGA-PAAD cohorts were acquired from the TCGA Pan-Cancer Atlas via cBioPortal, while baseline expression profiles for human cancer cell lines were retrieved from the Cancer Cell Line Encyclopedia (CCLE). A Macropinocytosis Score (MScore) was calculated for each sample by averaging the expression of a defined 72-gene signature. Patient cohorts were median-split into High and Low MScore groups, and overall survival was evaluated using the Kaplan-Meier estimator and log-rank test.

Differential expression between High and Low MScore PAAD tumors was performed to assess the molecular crosstalk of SASP-related proteases. To validate the signature *in vitro*, baseline cell line MScores were correlated with experimental growth inhibition following IPA3 treatment (100 µM) using the Pearson correlation coefficient. Data processing and visualizations were executed in Python.

## Statistical analysis

All experiments were performed with 4-6 technical replicates for each condition. The average and standard deviation were obtained and then the student’s t-test was used for comparisons between two groups and non-parametric Anova for comparison between multiple groups. The P-values used for statistical significance were as follows: P<0.05 was considered statistically significant with following notation for statistical significance ****<0.0001, ***<0.001, **<0.01, *<0.05. All statistical analysis was performed using Graphpad Prism v10.

## Results

### eATP increases iATP in multiple cancer cell lines *in vitro* – short-time study

We first investigated intracellular ATP (iATP) accumulation as phenotypic change associated with extracellular ATP (eATP) quantified intracellular ATP (iATP) following extracellular ATP (eATP) exposure (0.25–0.75 mM) over 30–120 min across multiple cancer cell lines representing distinct tissue origins. eATP significantly elevated iATP relative to untreated controls in all cell lines tested. In most lines, responses were comparable between 0.5 and 0.75 mM, indicating that 0.5 mM eATP was sufficient to achieve near maximal iATP elevation. However, time-course patterns differed by cell line (e.g., early plateau for Skhep1 and HOP92 vs progressive accumulation for HT29, Panc1, MCF7 and H1299), consistent with our hypothesis of heterogeneous uptake capacity (Figure 1A-F). These data demonstrate that eATP rapidly and robustly elevates iATP across diverse cancer cell types *in vitro*.

**Figure 1.**
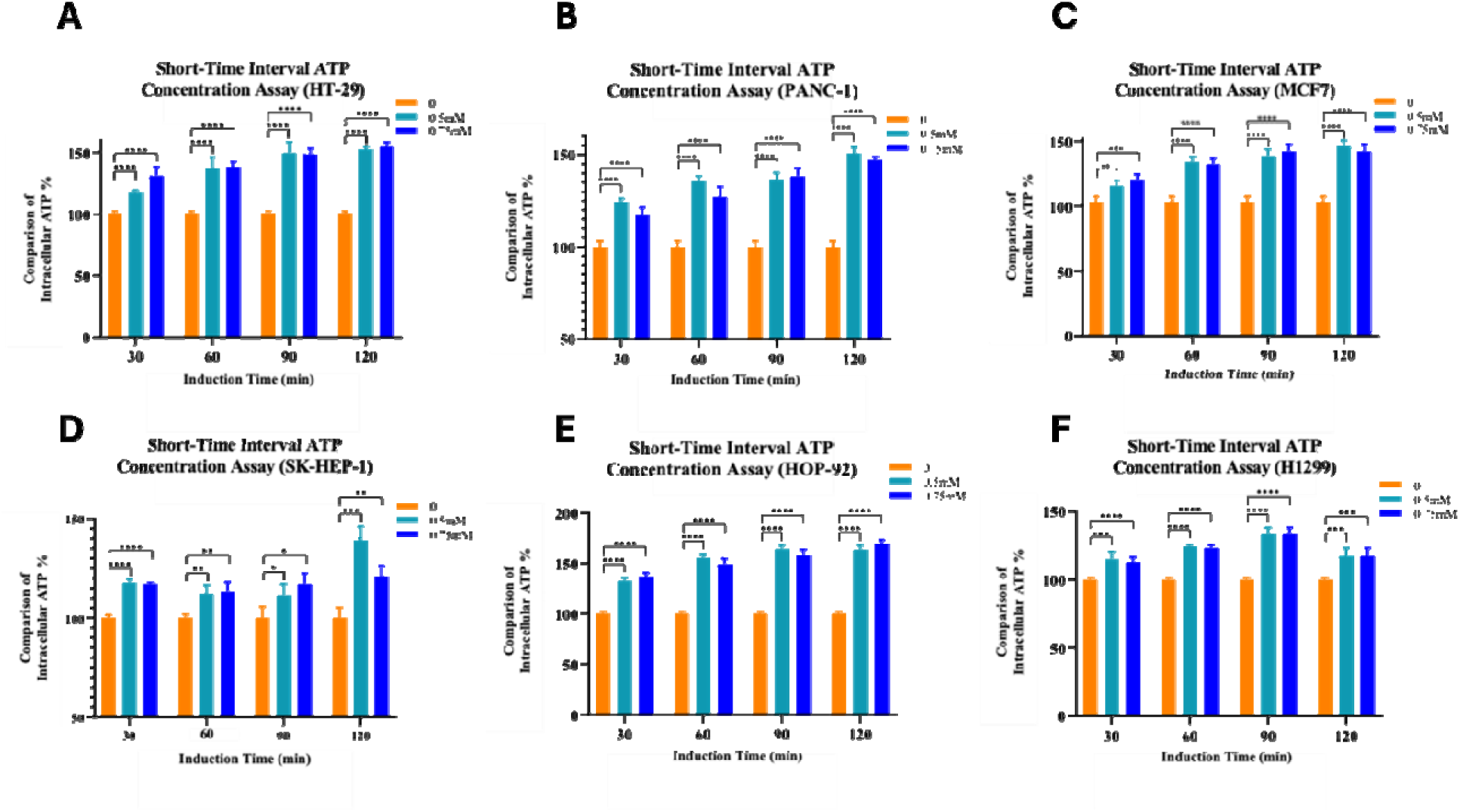
Short-time interval intracellular ATP uptake in diverse cancer cell lines. **(A–F)** Intracellular ATP (iATP) concentration assays at short-time induction intervals (30-120 minutes) in seven cancer cell lines based on 0, 0.5 and 0.75 mM eATP concentration: **HT29 (A), PANC1 (B), MCF7 (C), SK-HEP1 (D), HOP-92 (E), H1299 (F)**. Data is representation for one set of biological replicates as mean ± SD for (n ≥ 5 technical replicates). One-way ANOVA with post-hoc multiple comparisons; *P < 0.05,**P < 0.01,***P < 0.001, ****P < 0.0001. Orange, cyan, and blue bars correspond to increasing ATP concentrations as shown.

### eATP elevates iATP in a cell line–dependent manner *in vitro* over longer incubation

We next examined whether prolonged eATP incubation (2-4 hours) further influenced iATP accumulation beyond 2 hours. Heterogeneity in iATP increase with distinct, cell line–dependent responses were observed again as in case of short time incubation (Figure 1). Some cell lines had sustained elevation (HT29 and H1299) while others plateaued (Panc1, HOP92, MCF7 and Skhep1). However, differences were observed between 0.25, 0.5 and 0.75 mM dosages at all three time points for almost all the cell lines. Overall, prolonged eATP exposure drives iATP elevation in a cell line–dependent manner, consistent with differences in macropinocytic/endocytic capacity and/or ATP handling (Figure 2A-F).

**Figure 2.**
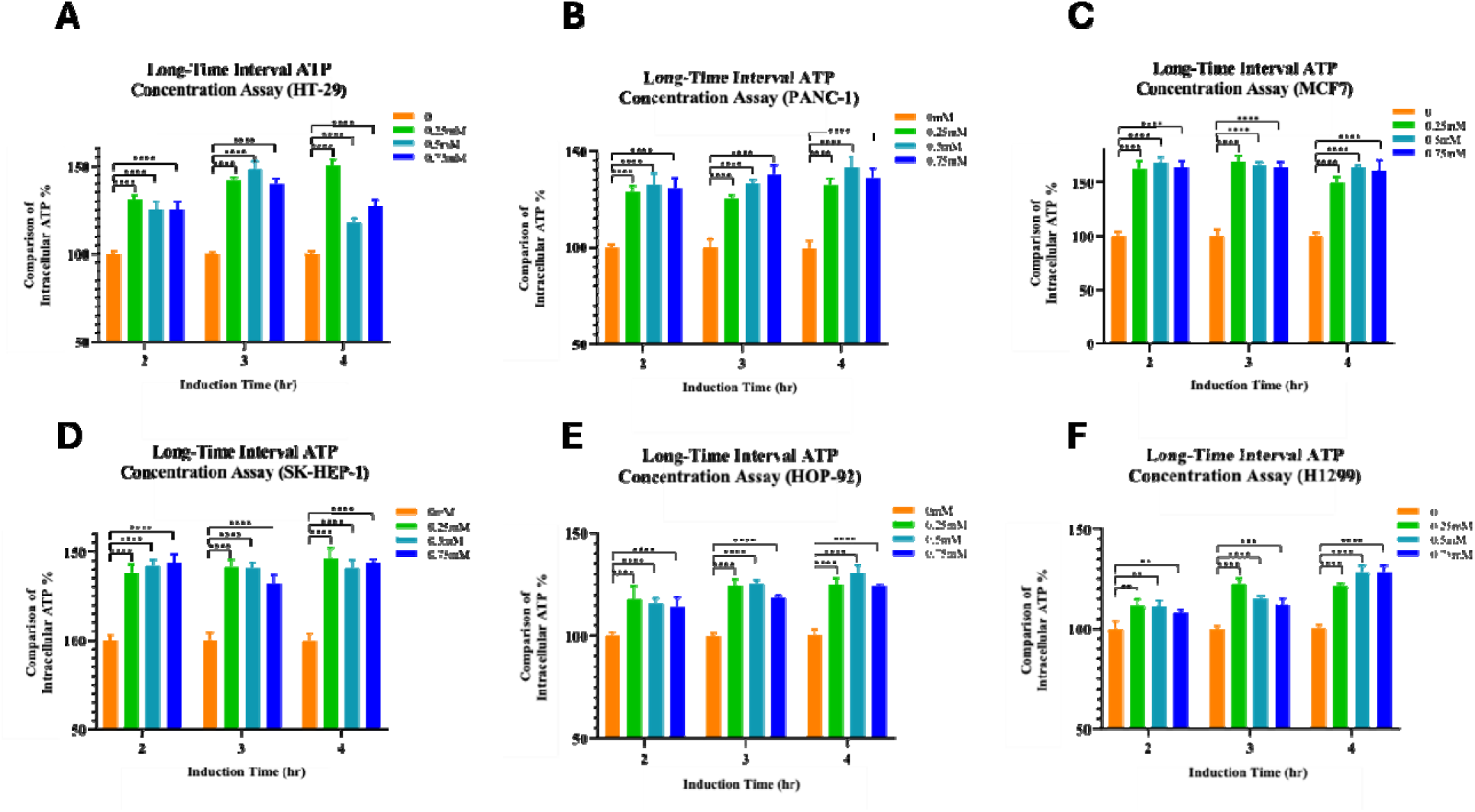
Long-time interval intracellular ATP uptake in diverse cancer cell lines. **(A–F)** Intracellular ATP (iATP) concentration assays at longer-time induction intervals (2-4 hours) in seven cancer cell lines based on 0, 0.25, 0.5 and 0.75 mM eATP concentration: **HT29 (A), PANC1 (B), MCF7 (C), SK-HEP1 (D), HOP-92 (E), H1299 (F)**. Data is representation for one set of biological replicates as mean ± SD for (n ≥ 5 technical replicates). One-way ANOVA with post-hoc multiple comparisons; *P < 0.05,**P < 0.01,***P < 0.001, ****P < 0.0001. Orange, green, cyan, and blue bars correspond to increasing ATP concentrations as shown.

### Macropinocytosis inhibitor IPA3 reduces eATP-induced iATP accumulation

Ww next tested whether the macropinocytosis inhibitor IPA3 could block eATP induced iATP increase across all cancer cell lines. Cells were treated with increasing concentrations of IPA3 (0-100 µM) for 30 minutes in the presence of 0.5 mM eATP. IPA3 reduced eATP-induced iATP accumulation in a dose-dependent manner in several cell lines, with variable sensitivity across cancer types. Some lines showed strong inhibition at lower doses (Panc1, Skhep1 and HOP92), whereas others required higher concentrations (MCF7, H1299 and A549) or displayed limited inhibition (HT29), supporting heterogeneity in macropinocytosis dependence. These results indicate that macropinocytosis contributes substantially to eATP-driven iATP elevation (Figure 3A-G).

**Figure 3.**
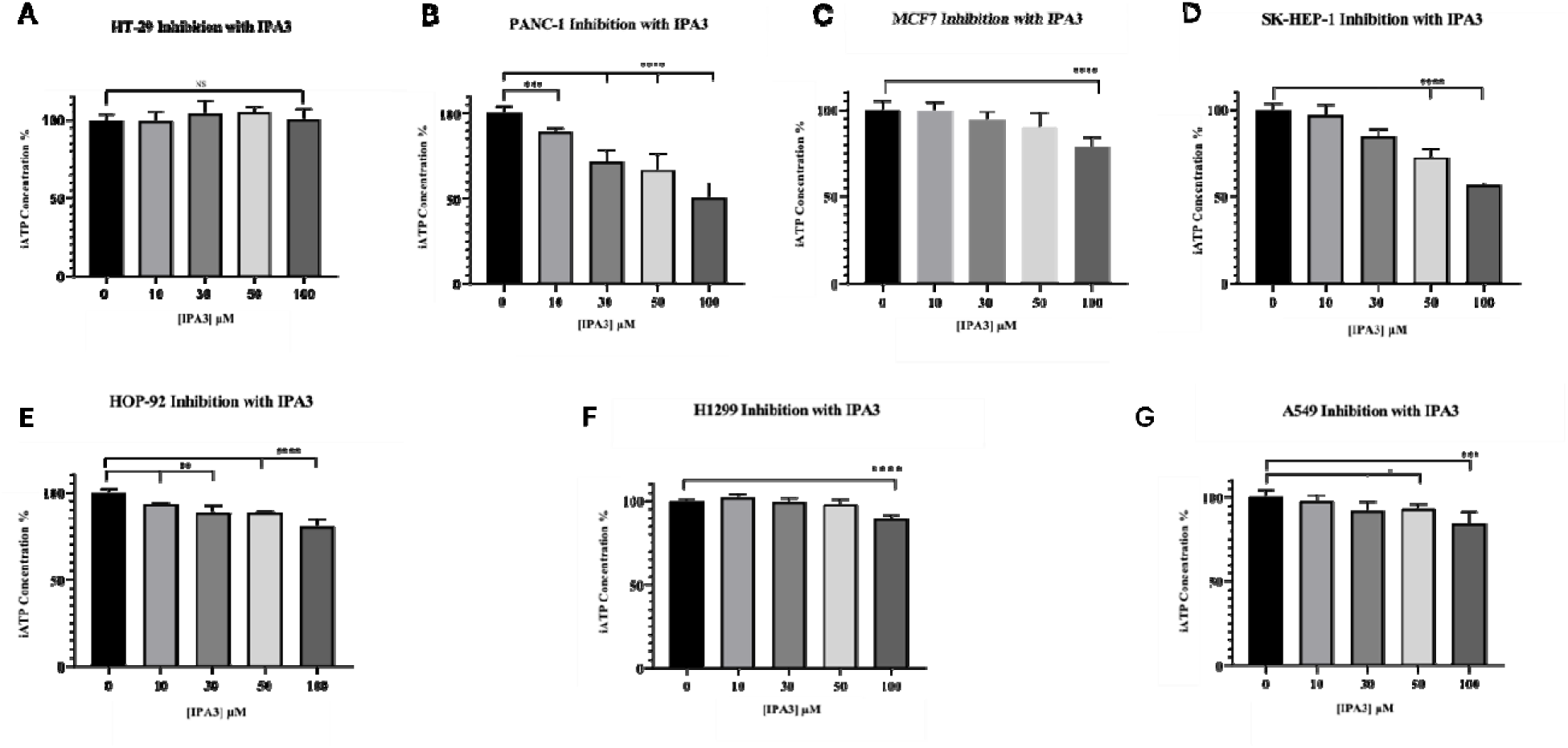
IPA3 inhibition of intracellular ATP accumulation across cancer cell lines. **(A–H)** Dose-dependent inhibition of intracellular ATP (iATP) different concentrations (0-100 µM) by IPA3 (PAK1 inhibitor) in eight cancer cell lines: **HT29 (A), PANC1 (B), MCF7 (C), SK-HEP1 (D), HOP-92 (E), H1299 (F), and A549 (G).** Cells were treated with increasing concentrations of IPA3 (µM) and 0.5 mM eATP for 30 minutes and subsequently iATP (%) was measured relative to untreated controls. Data is representation for one set of biological replicates as mean ± SD for (n ≥ 5 technical replicates). One-way ANOVA with post-hoc multiple comparisons; *P < 0.05,**P < 0.01,***P < 0.001, ****P < 0.0001; NS: non-significant.

### STC1 protein is key regulator of macropinocytosis in NSCLC *in vitro*

Given prior transcriptomic evidence that STC1 is upregulated following eATP exposure, we evaluated ATP uptake using A549 control and A549 STC1-KO cells. Short-time iATP elevation was similar between control and STC1-KO cells (Figure 4A-B), whereas longer exposure revealed significantly reduced iATP accumulation in STC1-KO cells relative to controls (Figure 4C-D). Consistent with this, fluorescence microscopy showed qualitatively reduced NHF-ATP/HMWFD colocalization in STC1-KO cells, while non-tumorigenic NL-20 cells displayed minimal uptake (Figure 4E). These results identify STC1 as an important regulator of macropinocytic ATP internalization in NSCLC cells and validated both genetic and pharmacological inhibition of macropinocytosis.

**Figure 4.**
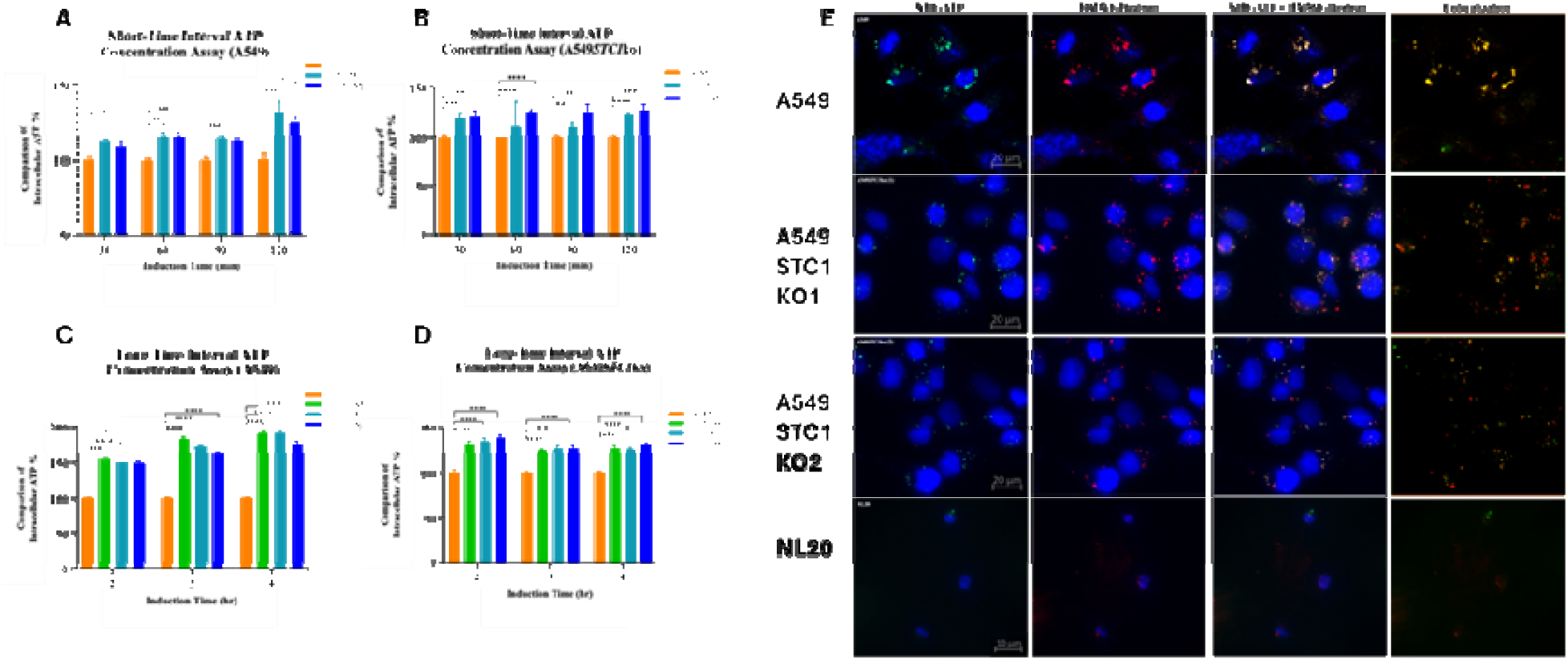
STC1 loss reduces macropinocytosis-driven intracellular ATP accumulation in A549 cells. **(A–B) Short-time interval ATP concentration assay (minutes).** Intracellular ATP (iATP, % of control) for A549 (A) and A549STC1KO (B) cells measured at the indicated induction times between 30-120 minutes. **(C–D) Long-time interval ATP concentration assay (hours).** iATP (%) for A549 WT (C) and STC1-KO (D) across induction times between 2-4 hours. **(E) Immunofluorescence/confocal imaging of macropinocytosis.** A549 WT, A549 STC1-KO (two independent clones), and NL20 control cells were stained with NHF-ATP (green; ATP analog), High molecular weight dextran (red-macropinosome marker), and DAPI (blue; cell nuclei). Merged images show colocalization (yellow). STC1-KO cells display visibly reduced FITC-dextran uptake relative to WT based on gene expression reduction, while NL20 shows minimal uptake. Scale bars: 20 µm (A549 and STC1-KO), 10 µm (NL20). Data is representation for one set of biological replicates as mean ± SD for (n ≥ 5 technical replicates). Colors denote the eATP dose as indicated in the in-plot legend. One-way ANOVA with post-hoc multiple-comparisons; *P < 0.05,**P < 0.01,***P < 0.001, ****P < 0.0001.

### Fluorescence microscopy confirmed NHF-ATP colocalization with HMWFD in cancer cells in vitro *and* in vivo

To validate the macropinocytosis mechanism underlying eATP uptake, we performed confocal fluorescence microscopy using high molecular weight fluorescein–dextran (HMWFD, red) as a fluid-phase marker of macropinosomes and a non-hydrolyzable fluorescent ATP analog (NHF-ATP, green) as a surrogate for ATP internalization. Colocalization of NHF-ATP with HMWFD was observed as yellow puncta in merged images, confirming macropinocytosis-driven uptake. We visualized ATP uptake using a non-hydrolyzable fluorescent ATP analog (NHF-ATP) together with high-molecular-weight fluorescent dextran (HMWFD), a fluid-phase marker of macropinosomes. Across multiple cancer cell lines, NHF-ATP colocalized with HMWFD as punctate intracellular structures, consistent with macropinocytic internalization (Supplementary Figure S1A-G). The extent of uptake and colocalization varied across lines, suggesting cell type–dependent differences in macropinosome formation, vesicle size, and the proportion of cells exhibiting uptake. Breast cancer MCF7 cells (Figure S1C) internalized NHF-ATP and HMWFD, but colocalization was minimal, suggesting alternative endocytic mechanisms may also contribute. NSCLC H1299 cells (Figure S1E-F) displayed strong colocalization, with more prominent macropinosome formation than HOP92. These data provide direct imaging evidence that macropinocytosis is a prevalent mechanism of ATP internalization in vitro. After confirming macropinocytosis *in vitro*, we wanted to test *in vivo* conditions. We injected NHF-ATP, dextran or both in mice (N=1) for 15 minutes and then mice were sacrificed and tumors were collected, fixed in OCT and then tissue sections were checked with fluorescence microscopy. NHF-ATP and dextran colocalized to varying degrees across tumor models, supporting in vivo ATP internalization with notable heterogeneity in uptake and vesicle morphology (Figure 5A-H).

**Figure 5.**
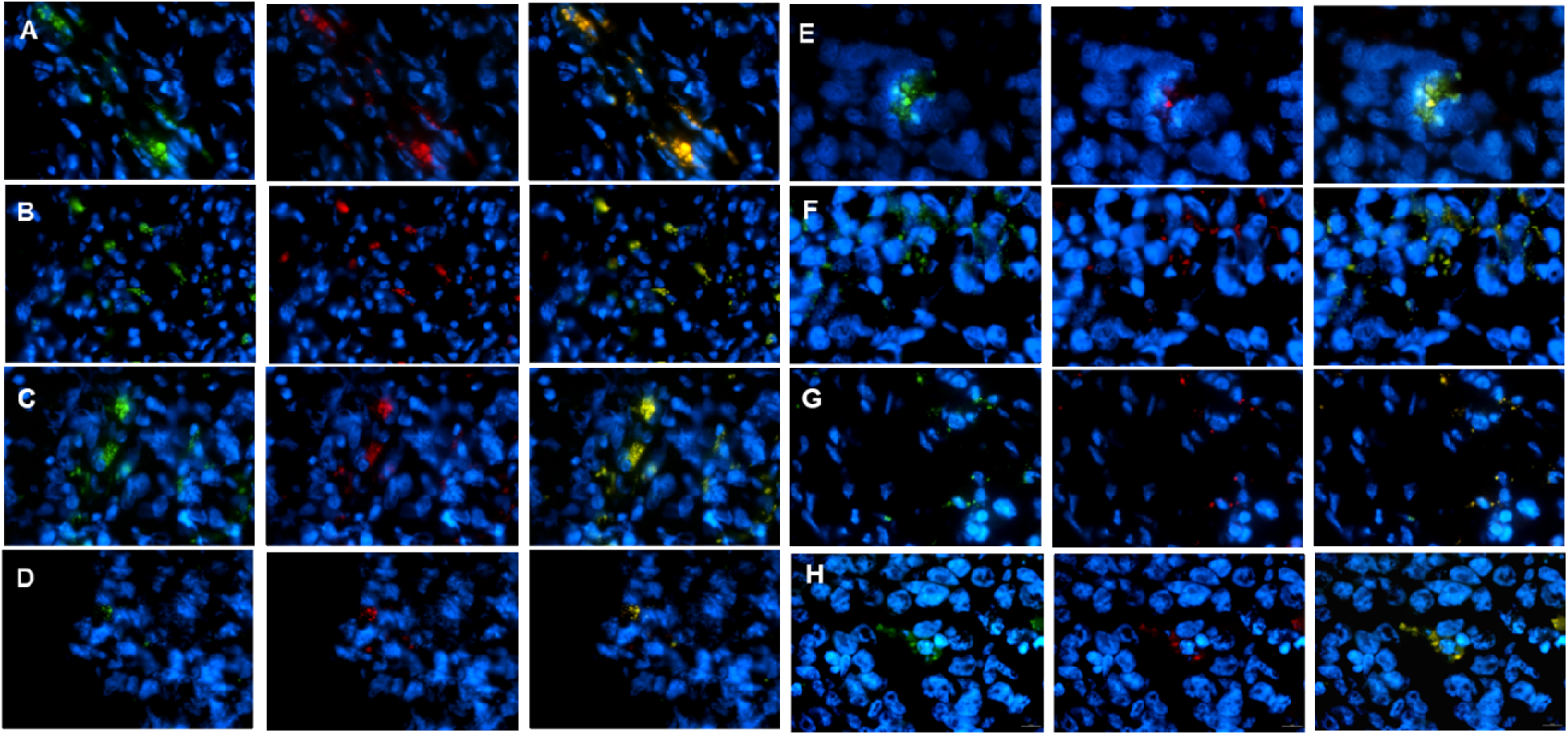
NHF-ATP is internalized and colocalizes with HMWFD *in vivo* in cancer cells. Tumor xenografts were established by subcutaneous injection of 1×10 –5×10 cells into the right flank of 3–4-week-old male NU/J nude mice. When tumors reached ∼200–500 mm³, tumors were injected intratumorally with DMEM control, HMWFD tracer, or HMWFD plus NHF-ATP (100 μM) in a total volume of 50 μL using 27G needles. Tumors were harvested within ∼7–8 minutes post-injection, embedded in OCT, and cryosectioned (10 μm). Sections were ethanol-fixed, PBS-washed, and mounted with antifade medium containing DAPI. Images were acquired using a Nikon NiU epifluorescence microscope with identical exposure settings across groups and processed using Nikon NIS-Elements software. **A-H.** NHF-ATP and HMWFD colocalize in **HT29 (A), PANC1 (B), MCF7 (C), SK-HEP1 (D), HOP-92 (E), H1299 (F), and A549 (G) and A375 (H).** respectively. Color: Blue:DAPI (nuclei), Red: HMWFD, Green: NHF-ATP, Yellow: Co-localization. Scale bar: 10 µm, 100x magnification.

A375 melanoma cells exhibited colocalization higher than other cancer cells (Figure 5H). HOP92 cells had more NHF-ATP rather dextran and although colocalization, ATP internalization was governed by other endocytic mechanisms as well (Figure 5E). H1299 cells had lower colocalization than A375 but macropinosomes were larger in size than A549 (Figure 5D). MCF7 macropinosomes had large size but smaller than HT29 and complete colocalization but some NHF-ATP was internalized in smaller endosomes were as well similar to what was observed *in vitro* (Figure 5C). Skhep1 cells have less degree of NHF-ATP internalization but complete colocalization (Figure 5D). Overall, some cell lines had similar internalization compared to *in vitro* studies again highlighting heterogeneity in this phenomenon.

### Clinical Validation and Prognostic Impact of Macropinocytic Capacity

To evaluate the clinical relevance of our experimental findings, we utilized a 72-gene macropinocytosis signature to calculate a transcriptomic score (MScore) across clinical cohorts from the TCGA Pan-Cancer Atlas. We first correlated our in vitro IPA3 inhibition data with CCLE cell line Mscore (Supplementary Figure S2) with Pearson R value of 0.63. Although modest this data can beomce more significant with the addition of IPA3 inhibition data from more cancer cell lines. In the lung adenocarcinoma (LUAD) cohort, patients were stratified into High and Low MScore groups based on a median split. Kaplan-Meier survival analysis revealed that patients with a High MScore exhibited significantly reduced overall survival compared to those in the Low MScore group (p = 0.009, Log-rank test; Figure 6A). This suggests that the eATP-driven scavenging mechanism identified in our *in vitro* A549 models correlates with a more aggressive clinical phenotype and poor prognosis in lung cancer.

**Figure 6:**
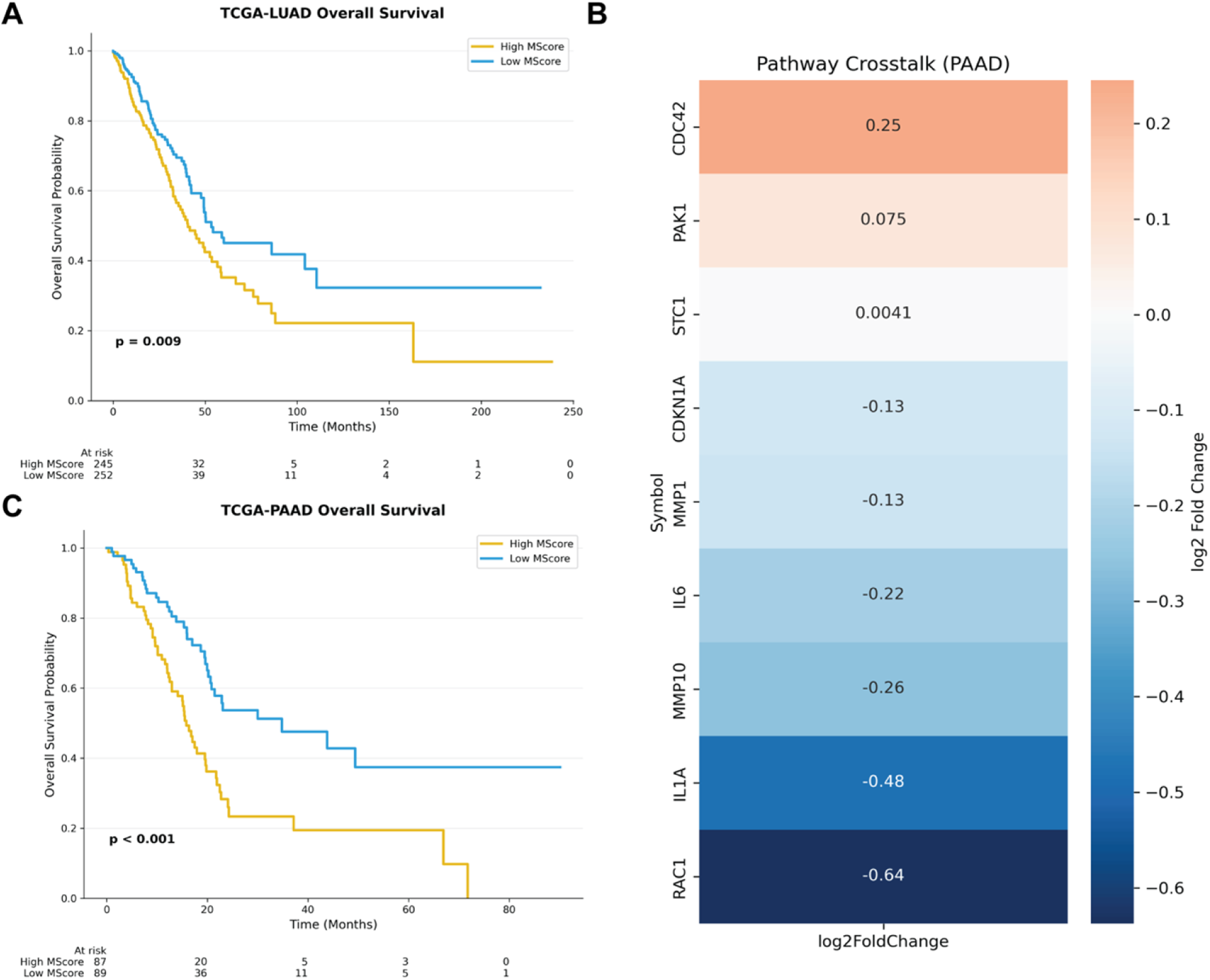
Clinical Validation and Prognostic Impact of Macropinocytic Capacity. **A.** Kaplan-Meier overall survival curve for the TCGA-LUAD cohort stratified by Macropinocytosis Score (MScore). Patients with a High MScore (in yellow) show a worse survival trend compared to those with a Low MScore (in blue; p = 0.009) **B.** Heatmap representing the log2 fold change of macropinocytosis and SASP-related genes in High-MScore vs. Low-MScore PAAD tumors. **C.** Kaplan-Meier overall survival curve for the TCGA-PAAD cohort stratified by MScore. The 177 patients in the pancreatic cancer cohort show a distinct split in survival based on their metabolic scavenging score (p < 0.001), highlighting macropinocytosis as a baseline survival requirement in the nutrient-poor pancreatic microenvironment. (

Furthermore, the most striking evidence of our dual-pathway model emerged from the transcriptomic crosstalk analysis. Differential expression analysis between High and Low MScore PAAD tumors revealed a significant upregulation of key senescence-associated secretory phenotype (SASP) factors in high-scavenging tumors. Specifically, macropinocytosis hub genes, such as CDC42, were positively correlated with the expression of pro-inflammatory proteases, including MMP10, MMP1, and MMP3 (Figure 6B). We further extended this analysis to pancreatic adenocarcinoma (PAAD), a malignancy characterized by an extremely nutrient-poor microenvironment where macropinocytosis is a recognized survival strategy. The PAAD cohort showed a highly significant and distinct metabolic survival profile (p < 0.001, Figure 6C). These clinical data suggest that high macropinocytic capacity provides the metabolic fuel necessary to maintain a robust SASP, effectively bridging our mechanistic discovery of eATP-driven ‘cellular drinking’ with the aggressive secretome observed in human pancreatic and lung malignancies.

### Temporal variation in senescence-associated gene expression and metabolite profiles following eATP treatment in A549 cells between 2 to 6 hours

We previously found eATP induces drug resistance, EMT and CSC phenotypic in cancer cells so we wanted to investigate if it could also induce senescence associated secretory phenotype (SASP) and so transcriptomics as well as metabolomics data ^6,7,33^ was reanalyzed to look for senescence formation related biomarker genes and metabolites. In the RNAseq data, multiple senescence-related genes and SASP-associated cytokine/chemokine and matrix-remodeling transcripts exhibited induction patterns consistent with stress-response remodeling ^3,5,7,34,35^. eATP treatment showed strong temporal concordance between early (2 hours) and late time points (6 hours), with most genes maintaining directional regulation and clustering near the diagonal (Pearson R = 0.87, P < 10 ¹), indicating a relatively stable transcriptional response program over this interval (Figure 7A). Only a limited subset of genes exhibited direction switching between time points. In contrast, TGF-β treatment displayed weak temporal concordance (Pearson R = 0.08, P = 0.68; NS), with broader dispersion and increased quadrant switching, consistent with greater transcriptional reprogramming between early and later responses (Figure 7B). Under eATP treatment, several metabolites showed substantial magnitude shifts between time points, including NAD , Gal-1-P, and fumarate, while others exhibited modest or opposite-direction changes, indicating heterogeneous metabolic remodeling rather than uniform temporal scaling (Figure 7C). TGF-β treatment produced a similarly dynamic metabolic pattern, with multiple intermediates in glycolytic and nucleotide metabolism pathways showing time-dependent divergence (Figure 7D). Consistent with the weak metabolite time-point correlation, these paired comparisons highlight directional and magnitude changes. Together, these results indicate that eATP induces a comparatively stable transcriptional program over time while metabolite levels undergo more heterogeneous remodeling,

**Figure 7.**
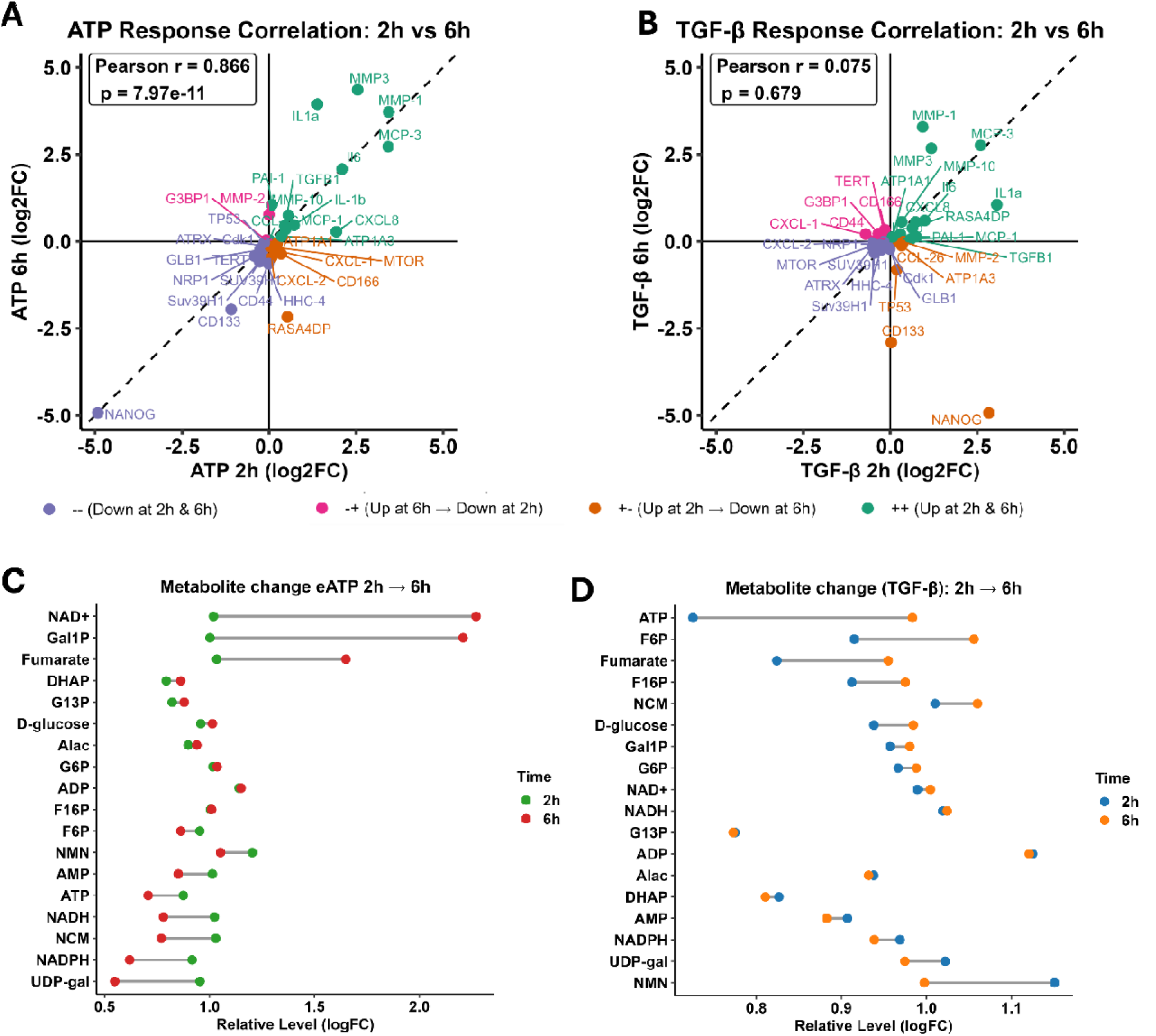
Transcriptomic and metabolomic analyses of A549 cells treated by eATP. RNA sequencing and metabolomics analyses datasets were used from previous study ^33^. A549 cells were treated with either 0.5 mM ATP (A) or 10 ng/mL TGF-β (T) for various times and then analyzed. Set of 34 genes and 18 metabolites belonging to senescence were selected (Supplementary data 1). **(A-B)** Scatter correlation plot of RNA-seq log2 fold-change values for senescence related genes at 2 h versus 6 h following eATP (**A**) and TGF-β (**B**) treatment. Each point represents one gene. The dashed diagonal indicates equal response at both time points. Solid horizontal and vertical lines denote zero fold-change. Points are color-coded by temporal directionality: ++ (up at 2 h and 6 h), −− (down at 2 h and 6 h), +− (up at 2 h → down at 6 h), and −+ (down at 2 h → up at 6 h). Pearson correlation coefficient and p-value are shown. **(C-D)** Paired dumbbell plot showing metabolite abundance changes between 2 h and 6 h after eATP (**C**) and TGF-β (**D**) treatment. Each metabolite is represented by a connected pair of points (green = 2 h, red = 6 h for eATP and blue = 2 h, orange = 6 h for TGF-β). Lines connect matched measurements, highlighting direction and magnitude of metabolic shifts over time. Several metabolites exhibit time-dependent remodeling rather than stable directional change for eATP, consistent with dynamic metabolic adaptation.

### eATP induced dose- and time-dependent expression of SA-**β**-Gal activity thereby inducing senescence in NSCLC cell lines

After confirming variations in genes associated with senescence, we investigated experimentally if eATP could induce it in lung cancer cells. Figure 8A schematically represents the design of the entire senescence study. SASP gene SA-β-Gal was increased in many cancer types including LUAD (Figure 8B) ^36,37^. Using three complementary SA-β-Gal assays and flow cytometry, we found that eATP induced a time-dependent senescence signal in A549 cells. eATP also induced time-dependent expression of SA-β-Gal (Figure 8C-E). eATP-induced SA-β-Gal appeared relatively fast, the increase could be observed in the first two hours after the treatment.

**Figure 8.**
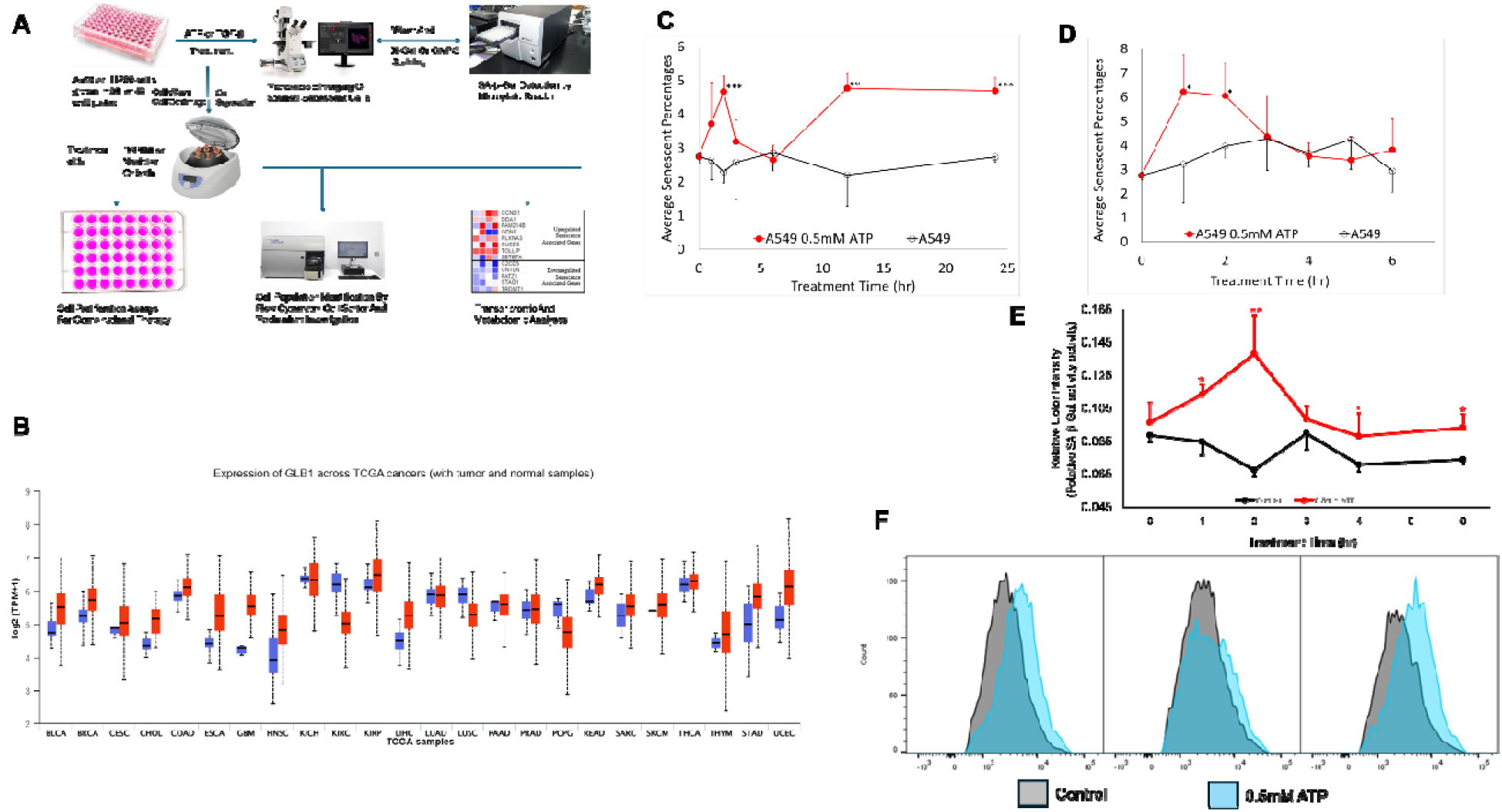
eATP induced SA-β-Gal expression in NSCLC A549 cells. Chromogenic assays of eATP-induced SA-β-Gal expression in NSCLC A549 cells were performed. The time-dependence response of SA-β-Gal expression to 0.5 mM eATP was measured by manual counting, plate reader, and flow cytometry. Each assay was repeated for at least 3 times, and the same experimental conditions were repeated 3-6 times (N= 3-6). *P < 0.05. Unpaired t-test was performed using **A**. Schematic representation of the study design. **B**. Higher SA-β-Gal gene expression level is correlated with tumor compared to normal tissues. **C**. Time-dependent SA-β-Gal activity of A549 cells treated with 0.5 mM eATP (from 0 – 24 hours). **D**. Shorter but more detailed time-dependent SA-β-Gal activity of A549 cells treated with 0.5 mM eATP (from 0 – 6 hours). **E**. Manual reading of A549 senescence cells after eATP treatment. **F**. A subpopulation of SA-β-Gal expressing A549 cells was induced by 0.5 mM eATP and detected by flow cytometry.

Interestingly, it declined at 3-5 hours and restored after 6 hours and persisted for at least 24 hours (Figure 8 C-E). Additionally, X-Gal-based plate reading assay (Figure 8F) and flow cytometry also showed qualitatively similar results (Figure 8G). Because SA-β-Gal is one of the most prominent biomarkers for senescence, we concluded from this study that eATP induces senescence in human NSCLC A549 cells.

Next, we investigated if eATP induces senescence in another NSCLC cell line, H1299. eATP also induced expression of SA-β-Gal in H1299 NSCLC cells (Supplementary Figure S3A), Further senescence induction validation was observed using ONPG-based plate reading assay (Supplementary Figure S3B). These assay results indicate that eATP-induced senescence in NSCLC cells might be a common phenomenon.

### Purinergic receptor signaling mediates eATP-induced senescence independently of macropinocytosis

We then investigated the downstream signaling mechanism of eATP induce senescence. Flow cytometry analysis of A549 cells treated with either eATP (blue) or eATP plus a PR signaling inhibitor oATP (purple) showed a time-dependent shift with reduced senescent cell population selectively for oATP treated samples (Figure 9A) for 2 (panel A) but not for 6 hours (panel B) consistent with timings obtained in previous assays. However, IPA3 did not show such decrease shift as both 2 (Panel A) and 6 hours (panel B) between eATP (blue) or eATP+IPA3 (green) (Figure 9B). These results indicate that macropinocytosis does not play a significant role in the senescence induction by eATP. Finally, we summarized the mechanism of eATP induced senescence (Figure 10C).

**Figure 9.**
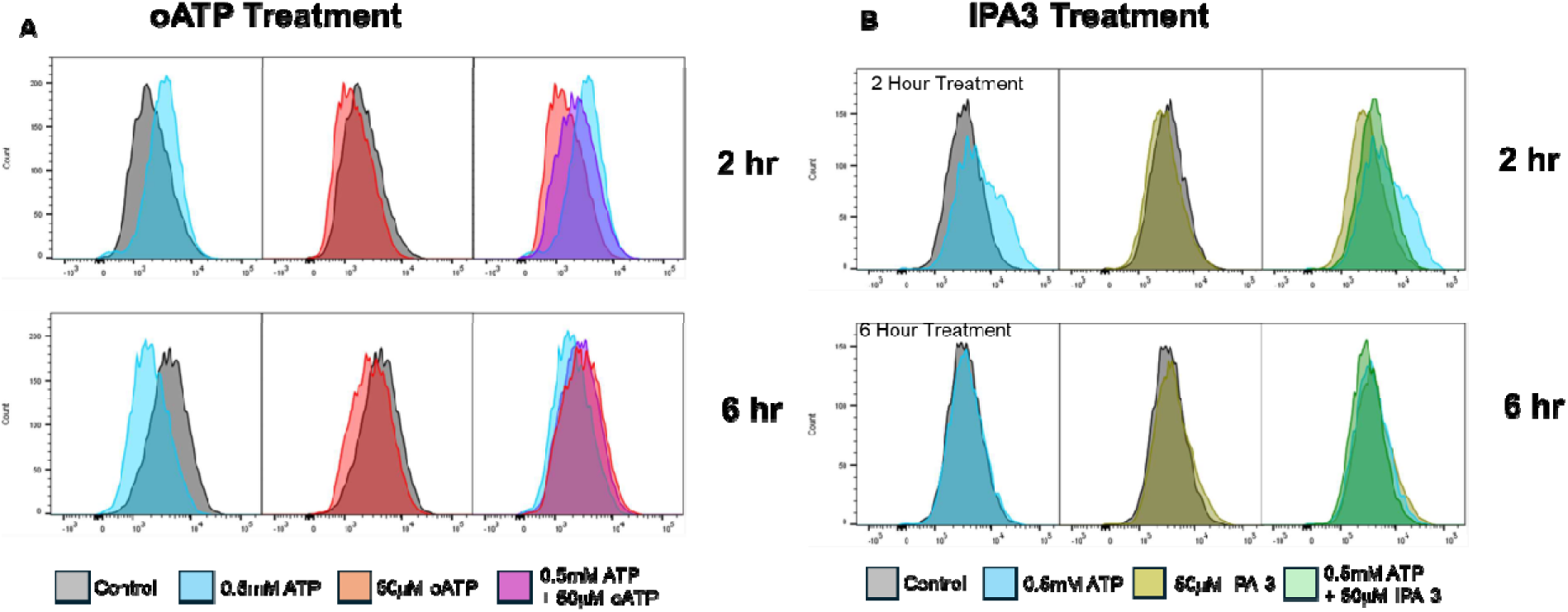
eATP induces senescence primarily via signaling of purinergic receptor P2X7. A549 cells were treated with either 0.5 mM eATP or 50 mM oATP or 50 mM IPA-3 for various hours. After the treatments, the treated cells were analyzed by flow cytometry. **A**. PR (P2X7) inhibitor oATP significantly reduced SA-β-Gal activity in A549 cells. **B**. Macropinocytosis inhibitor IPA-3 did not reduce SA-β-Gal activity.

## Discussion

Cancer cells rely on opportunistic nutrient acquisition to sustain their elevated metabolic demands, and macropinocytosis has emerged as a major pathway supporting nutrient scavenging, metabolic flexibility, and therapy resistance ^11,38–40^. Macropinocytosis is also an escape mechanism by which cancer cells can overcome nutrient deprivation by uptake of glutamine or glucose as well as a mechanism increasing drug resistance ^6,15,30,41–43^. In this study, we extend our previous findings and demonstrate that extracellular ATP (eATP), which is present at high concentrations in the tumor microenvironment, is broadly internalized across multiple cancer types through macropinocytosis both in vitro and in vivo. This uptake leads to rapid elevation of intracellular ATP (iATP) levels, with kinetics and magnitude varying across cell lines, consistent with heterogeneous macropinocytic capacity (Figure 1-2).

Pharmacological inhibition experiments supported macropinocytosis as a major contributor to eATP-driven iATP elevation. The macropinocytosis inhibitor IPA3 reduced iATP accumulation in most tested cancer cell lines, although sensitivity varied, suggesting that alternative endocytic routes or differential pathway activation may contribute in some contexts (Figure 3). Genetic evidence further supports this mechanism, as STC1 loss reduced long-duration ATP accumulation and vesicular colocalization, identifying STC1 as an important regulator of sustained macropinocytic ATP uptake. In contrast, non-tumorigenic lung cells showed minimal uptake, reinforcing that high macropinocytic activity is predominantly a cancer-associated phenotype (Figure 4). Imaging with fluorescent ATP analog and dextran confirmed vesicular colocalization consistent with macropinosome-mediated uptake both *in vitro* and *in vivo* (Figure 5, supplementary Figure S1) in all the cancer cell lines tested indicating that this process is pan-cancer phenomenon.

Transcriptomic scoring using a macropinocytosis gene signature (MScore) correlated with experimental uptake phenotypes and showed prognostic associations in selected cancer cohorts, suggesting that macropinocytosis capacity may be partially predictable from gene expression patterns, although larger datasets are required for robust modeling (Figure 6). Having demonstrated that eATP uptake through macropinocytosis elevates intracellular ATP and reshapes cellular metabolism, we asked whether eATP exposure also engages stress-response programs independent of nutrient acquisition. Because extracellular ATP is a known purinergic signaling ligand and tumor microenvironment stress signal, we reanalyzed our prior RNA-seq and metabolomics datasets to test for activation of senescence-associated pathways ^33^. The observed upregulation of CDC42 in High-MScore clinical samples aligns with our *in vitro* data identifying it as a master regulator of vesicle closure during macropinocytosis. We propose that the STC1-CDC42 axis facilitates the transition to a high-scavenging phenotype, which in turn provides the metabolic flexibility required for the cell to escape the growth-inhibitory effects of senescence while retaining the potent, pro-tumorigenic signaling of the SASP. We observed consistent modulation of multiple senescence-related genes ^6^ together with metabolite shifts linked to redox and nucleotide metabolism (Figure 7), supporting the interpretation that eATP triggers both metabolic adaptation and senescence-associated cellular programs.

Beyond metabolic effects, we showed that eATP also induces a senescence phenotype in NSCLC cells, supported by multiple SA-β-Gal assays and flow cytometry (Figure 8, Supplementary Figure S3). Mechanistically, this effect is primarily mediated by purinergic receptor signaling, particularly P2X7, as pharmacologic PR inhibition markedly reduced senescence induction. In contrast, macropinocytosis inhibition with IPA3 had minimal effect on the senescence readout (Figure 9). These data distinguish two parallel eATP response pathways: macropinocytosis-dependent ATP internalization that supports energy-intensive phenotypes such as EMT, stemness, and drug resistance, and purinergic receptor signaling that drives senescence largely independent of ATP uptake. This pathway separation helps explain how eATP can trigger distinct biological programs with different energetic requirements. The senescence response showed rapid and dynamic kinetics, with early SA-β-Gal induction followed by partial decline and later stabilization, paralleling early timing previously observed for eATP-induced EMT and cancer stem cell programs. This temporal alignment supports the concept that eATP functions as an early tumor microenvironment stress signal coordinating multiple adaptive phenotype.

Overall, our findings support a dual-pathway model in which tumor-level eATP acts both as a macropinocytic nutrient source and a purinergic signaling ligand. Targeting the macropinocytosis scavenging pathway, perhaps through inhibition of the P2X7 receptor or CDC42, represents a promising therapeutic strategy to starve the SASP and limit the clinical progression of aggressive lung and pancreatic malignancies.

## Patents

Not applicable.

## Supplementary Materials

Supplementary materials are attached separately.

## Author Contributions

Conceptualization, P.S. and X.C.; methodology, P.S., X.C.; software, N.S., R.W., C.N., S.A., Y.L., H.Z., J.S., S.P., S.C., D.R., S.B., N.J., P.S., and X.C.; validation, N.S., R.W., and P.S.; formal analysis, P.S. and X.C.; investigation, N.S., R.W., C.N., S.A., Y.L., H.Z., J.S., S.P., S.C., D.R., S.B., N.J., P.S., and X.C.; data curation, N.S., R.W., P.S.; writing— original draft preparation, P.S.; writing—review and editing, P.S., and X.C.; visualization, P.S.; supervision, P.S. and X.C.; project administration, P.S. and X.C.; funding acquisition, X.C. All authors have read and agreed to the published version of the manuscript.

## Institutional Review Board Statement

Not applicable.

## Informed Consent Statement

Not applicable.

## Data Availability Statement

Data are contained within the article. The RNAseq data is stored at NCBI GEO (Gene Expression Omnibus) with accession number: GSE160671.

## Acknowledgments

None

## Abbreviations

TCGA: Total Cancer genome Atlas
iATP: Intracellular ATP
eATP: Extracellular ATP
SASP: Senescence associated senescence phenotype

