## supplementary file for "Dual Pathways of Extracellular ATP Action in Cancer Cells: Purinergic Signaling–Driven Senescence and Macropinocytic ATP Internalization"

**Supplementary Files**

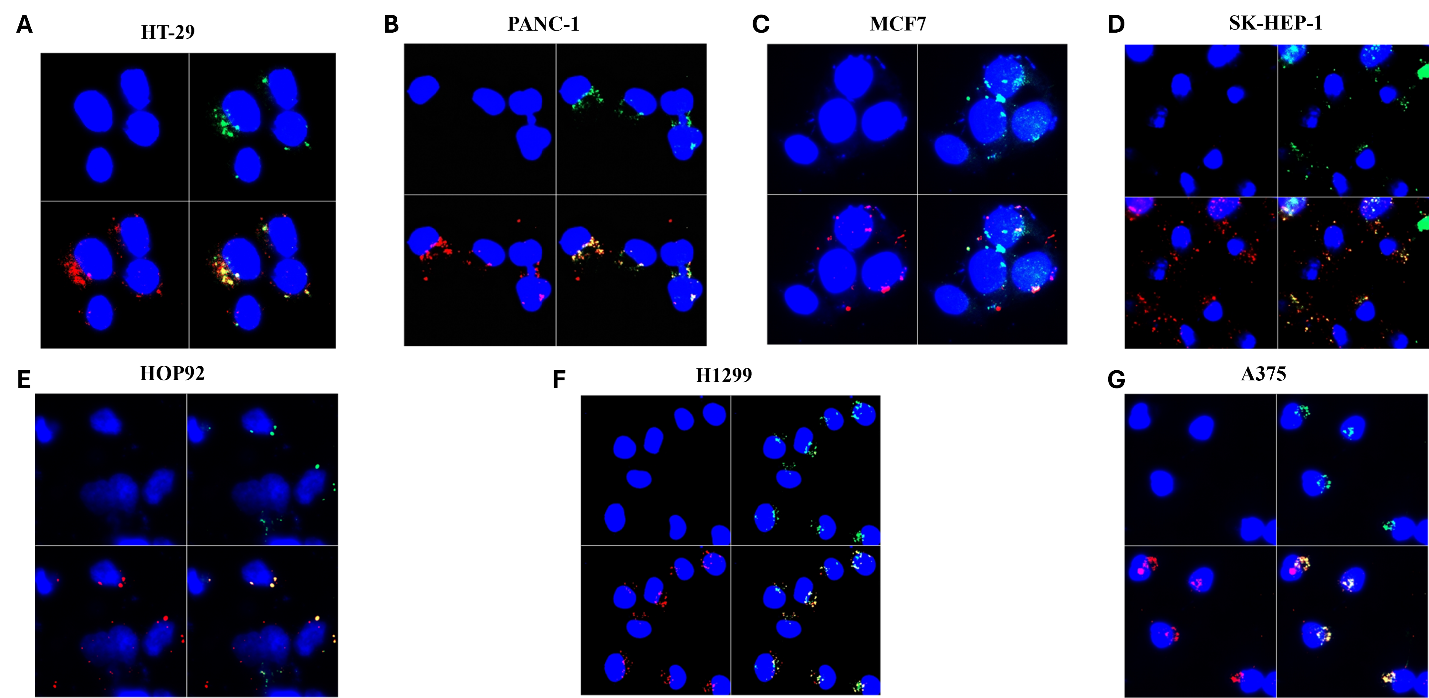

**Supplementary Figure S1. ATP internalization by macropinocytosis is ubiquitous phenomenon in cancer cells. (A–H)** Representative confocal immunofluorescence images of cancer cell lines showing macropinocytosis-driven uptake of a fluorescent ATP analog. Nuclei are stained with DAPI (blue), high molecular weight dextran is labeled in red (fluid-phase marker of macropinosomes), and NHF-ATP (non-hydrolyzable fluorescent ATP analog) is shown in green. Merged images demonstrate colocalization of dextran and NHF-ATP (yellow), confirming macropinocytosis-mediated internalization. **HT29 (A), PANC1 (B), MCF7 (C), SK-HEP1 (D), HOP-92 (E), H1299 (F), and A375 (G).** Differences in uptake intensity and vesicular localization highlight variation in macropinocytic activity between cancer cell lines. Each Figure has four panels; LT (left top) shows nuclei alone; RT (right top) shows nuclei alongwith NHF-ATP; LB (left bottom) shows nuclei along with HMWFD; RB (right bottom) co-localization of NHP-ATP and HMWFD. Scale bars: 20 µm.

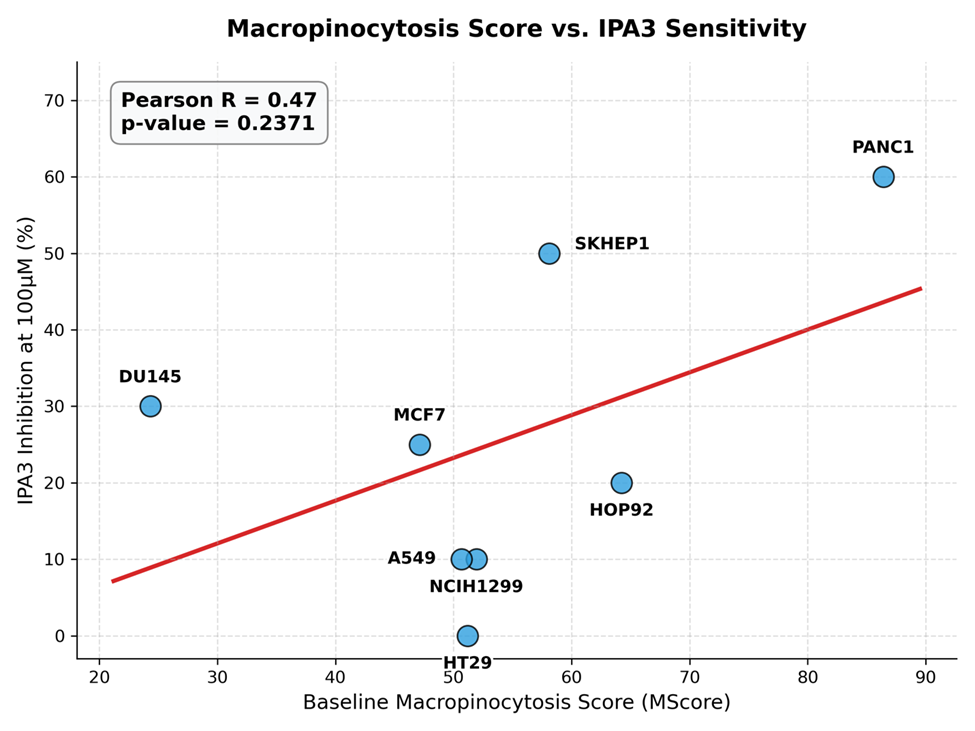

**Supplementary Figure S2.** **Correlation between baseline macropinocytosis capacity and pharmacological sensitivity.** Pearson correlation analysis comparing the transcriptomic Macropinocytosis Score (MScore) derived from CCLE baseline expression data and the experimentally determined cellular inhibition following treatment with the PAK1-inhibitor IPA3 (100 µM) across eight human cancer cell lines. High baseline MScore strongly correlates with increased sensitivity to IPA3-mediated inhibition, validating the dependency of high-scavenging cells on this specific pathway. CCLE gene expression data was collected from cbioportal.

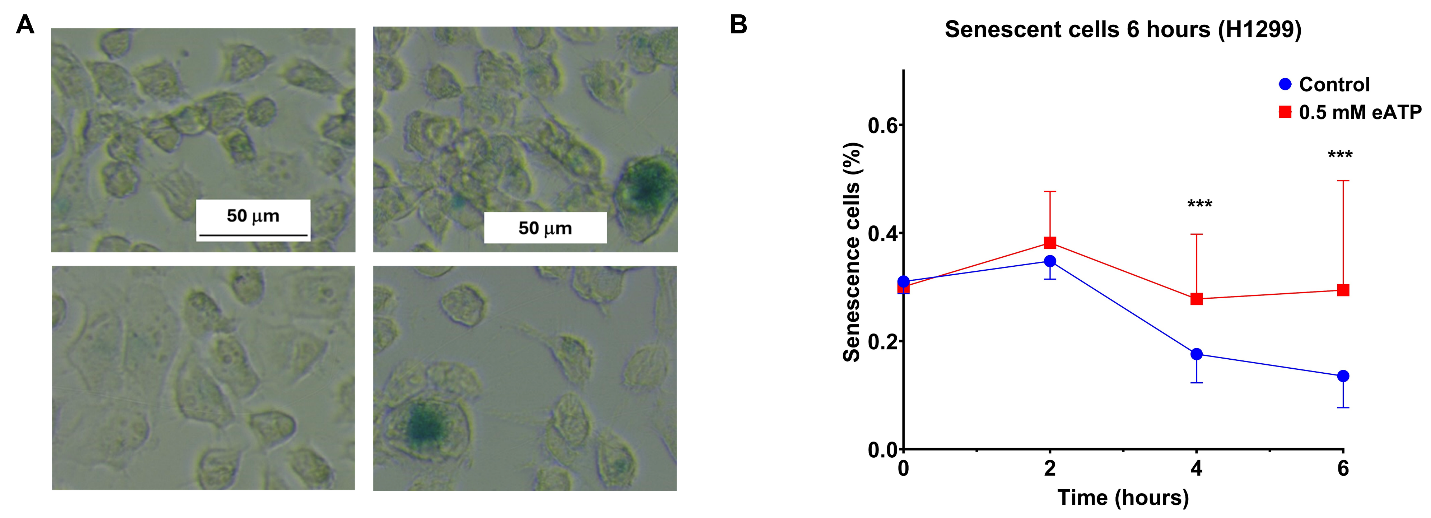

**Supplementary Figure S3.** eATP also induced SA- β -Gal activity in NSCLC H1299 cells. Senescence assays were performed in NSCLC H1299 cells. Each assay was repeated for at least 3 times, and the same experimental conditions were repeated 3-6 times (N= 3-6). *P < 0.05. **A**. eATP also induced SA- β -Gal activity (bluish) in NSCLC H1299 cells. **B**. eATP induced SA-β-Gal activity in H1299 cells as measured by plate reader.
